# A novel voltage clamp/dye uptake assay reveals saturable transport of molecules through CALHM1 and connexin channels

**DOI:** 10.1101/2020.02.15.950923

**Authors:** Pablo S. Gaete, Mauricio A. Lillo, William López, Yu Liu, Andrew L. Harris, Jorge E. Contreras

**Affiliations:** Department of Pharmacology, Physiology, and Neuroscience. New Jersey Medical School, Rutgers, The State University of New Jersey, Newark, NJ

**Keywords:** Hemichannel, connexin, CALHM1, permeability assay, saturable transport, large-pore channel

## Abstract

Channels that are permeable to small molecules such as ATP, in addition to atomic ions, are emerging as important regulators in health and disease. Nonetheless, mechanisms of molecular permeation and selectivity of these channels remain mostly unexplored due to the lack of quantitative methodologies. To address this need, we developed a novel two-electrode voltage clamp (TEVC)/dye uptake assay to examine the kinetics of molecular permeation of channels formed by human connexins (hCx), and the calcium homeostasis modulator (hCALHM1). We expressed hCx26, hCx30, and hCALHM1 individually in *Xenopus laevis* oocytes. To quantify the uptake of small molecular dyes through these channels, we developed a protocol that renders oocytes translucent – thereby amenable to optical detection techniques – without affecting the functional properties of the expressed channels. To control membrane potential and to determine functional channel expression accurately, dye uptake was evaluated in conjunction with TEVC. Using this methodology, we found that: (1) CALHM1 and Cx30 hemichannels display saturable transport of molecules that could be described by Michaelis-Menten kinetics, with apparent K_M_ and V_max_; (2) Kinetic parameters for molecular transport through CALHM1 are sensitive to voltage and extracellular calcium; (3) Significant transport of molecules occurs through CALHM1 when there are little or no ionic currents through the channels; (4) Cx mutations in the N-terminal region significantly affect kinetics of transport and permselectivity. Our results reveal that molecular flux through these channels has a rate-limiting step, that the kinetic parameters of molecular transport are sensitive to modulators of channel gating and that molecular transport and ionic currents can be differentially affected. Our methodology allows the analysis of how human mutations causing diseases affect kinetic properties and permselectivity of molecular signaling and enables the study of molecular mechanisms, including selectivity and saturability, associated with molecular transport in large-pore channels.

## INTRODUCTION

Large-pore channels are non-conventional membrane channels with a diverse amino acid sequence characterized by permeability to both atomic ions and small molecules. In vertebrates, examples of large-pore channels include those formed by connexins, pannexins, calcium homeostasis modulator (CALHM), and LRRC8 (SWELL channels). Although they share some common properties, such as membrane topology, pharmacological profiles and permeants, recent structural and functional evidence indicates that gating and selectivity mechanisms may vary for each channel.

Calcium homeostasis modulator protein 1 (CALHM1) is expressed mainly in the brain, and has been associated with the control of neuronal excitability and sensory perception (Dreses-Werringloer et al., 2008; Ma et al., 2012; Taruno et al., 2013). CALHM1 forms voltage-gated and Ca^2+^-sensitive channels with a wide pore (estimated to be > 14 Å diameter) that allow the non-selective transport of Ca^2+^, K^+^, Na^+^, monovalent anions and small molecules such as ATP and fluorescent dyes (Siebert et al., 2013; Ma et al., 2016; Syrjanen et al., 2020).

Connexins are the proteins that compose intercellular gap junction channels (GJCs) in vertebrates. GJCs are formed by the docking of two connexin hemichannels, each provided by adjacent cells. When hemichannels are undocked, they provide an autocrine/paracrine signaling pathway by a gated pathway between the intracellular compartment and the extracellular milieu (Saez et al., 2003). With 21 isoforms in humans, the connexin family represents the most diverse group of large-pore membrane channels. They are ubiquitously expressed in almost all human tissues. Based on structural data and permeability profiles, it is thought that the pore diameter of hemichannels is ~12-15 Å (Harris, 2007; Maeda et al., 2009; Sáez and Leybaert, 2014; Myers et al., 2018). Thus, connexin hemichannels have been described as conduits not only for atomic ions, but also for small biological molecules such as ATP, IP3, cAMP, glutamate, prostaglandins, and NAD^+^ (Harris, 2007).

The opening of large-pore channels, including connexin hemichannels and CALHM1 channels, is commonly evaluated by using conventional electrophysiological techniques and dye uptake/efflux assays, whose readouts are ionic currents and molecular permeability to fluorescent dyes, respectively. Nevertheless, recent evidence suggests that permeability to molecules cannot be extrapolated to atomic ion permeation and vice versa. The uptake of molecular dyes through connexin and pannexin channels has been described at resting membrane potentials where atomic ion currents are not detected (Contreras et al., 2003; Nielsen et al., 2019a). Furthermore, pharmacological blockade of connexins or pannexins have been shown to differentially affect atomic ion and molecular permeation (Nielsen et al., 2019a; Nielsen et al., 2019b). These data suggest that the mechanisms of transport for ions and molecules may be distinct. Yet, the most common and simple experimental strategy to evaluate the activity of these channels is through the uptake of small molecular fluorescent dyes such as ethidium, YO-PRO, propidium iodide, Lucifer Yellow and DAPI (Orellana et al., 2011; Siebert et al., 2013; Hansen et al., 2014b; Sáez and Leybaert, 2014; Gaete et al., 2019; Lillo et al., 2019). However, this methodology presents many experimental disadvantages, such as low temporal resolution and uncontrolled membrane potentials. Lack of normalization due to the inherent variability of protein expression and the number of functional channels located at the plasma membrane makes this methodology only a qualitative tool, insufficient to provide a basis for biophysical analysis of molecular permeation. To more rigorously study the biophysical properties of permeation in large-pore channels, an integrated approach is needed, capable of analyzing in the same preparation the kinetics of transport of molecules and atomic ions.

To address these issues, we developed a novel method that examines both ionic currents and permselectivity to fluorescent dyes simultaneously using Xenopus oocytes. By applying this methodology, we were able to distinguish novel and exciting biophysical properties in both connexin hemichannels and CALHM1 channels. The method reveals saturation of molecular flux and allows us to quantify kinetic parameters of molecular permeability (V_max_ and K_M_) at controlled membrane potential conditions. This methodology will also permit quantitative exploration of permeation of ions and molecules in other wide-pore channels including innexin, pannexin and SWELL channels.

## MATERIAL AND METHODS

### Collection of oocytes

Oocytes were collected from female *Xenopus laevis* according to the experimental protocol approved by the Institutional Animal Care and Use Committee (IACUC) at Rutgers University and conforming to the National Institutes of Health Guide for the Care and Use of Laboratory Animals. Briefly, frogs were anesthetized by immersion in a 0.35% ethyl 3-aminobenzoate methanesulfonate (MS-222) solution adjusted to pH 7.0-7.5. Subsequently, one ovarian lobe was excised to collect the oocytes. Oocytes were defolliculated using an OR-2 solution (composition in mM: 82.5 NaCl, 2.5 KCl, 1 MgCl_2_, 5 HEPES, adjusted to pH = 7.6) containing 0.3% collagenase type IA. Then, oocytes were stored at 16 °C in a ND96 solution (composition in mM: 96 NaCl, 2 KCl, 1 MgCl_2_, 1.8 CaCl_2_, 5 HEPES, adjusted to pH = 7.4) supplemented with streptomycin (50 μg/mL) plus penicillin (50 Units/mL).

### Molecular biology and channel expression

cDNA for human (h) CALHM1, hCx26, and hCx30 were synthesized by Epoach. cDNAs were amplified and PCR fragments were subcloned into pGEM-HA vector (Promega) for expression in Xenopus oocytes. The cDNAs were transcribed *in vitro* to cRNAs using the T7 Message Machine Kit (Ambion). Single mutations in hCx26 and hCx30 were produced using the QuickChange II Site-Directed Mutagenesis Kit (Agilent Technologies).

Healthy mature non-fertilized oocytes (stage IV-V) were individually microinjected with cRNA for CALHM1, Cx26, Cx26^N14K^, Cx30, and Cx30^G11R^ using the Nanoliter 2010 injection system (World Precision Instruments, Sarasota, FL, USA). Then, oocytes were stored separately in ND96 solution at 16 °C. After 1-2 days, oocytes were used for electrophysiology experiments or underwent a protocol to turn the animal pole translucent (see below). The amount of injected cRNA was 10-50 ng per oocyte.

Xenopus oocytes express endogenous Cx38, which could interfere with the analysis and interpretation of the acquired data (Ebihara, 1996). Therefore, to reduce the expression levels of Cx38, all the oocytes (including control oocytes not expressing exogenous proteins), were injected with an antisense oligonucleotide (0.5 μg/μL), using the sequence described previously (Ebihara, 1996).

### Preparation of translucent oocytes

Detection of fluorescence in intact Xenopus oocytes is challenging because of their high variable intrinsic autofluorescence and the high content of pigments and yolk granules that make difficult the detection of a fluorescent signal from the cytoplasm (Dumont, 1972; Lee and Bezanilla, 2019). To detect fluorescence emitted from the intracellular compartment, we rendered oocytes translucent. To achieve this, we optimized a centrifugation procedure previously reported by Iwao et al. (Iwao et al., 1997). Briefly, Eppendorf tubes containing 350 μL of 45% Ficoll® PM 400 were gently filled with 350 μL of ND96 solution with special care of not to mix the two solutions. Then, oocytes were individually immersed into the tubes so that the oocytes were retained at the interface between Ficoll and the ND96 solutions. In this interface, the animal pole of the oocyte faces up. The tubes were then centrifuged for 30 min at 1,700 g and 4° C. After centrifugation, oocytes were carefully collected and rinsed with ND96 to remove Ficoll. Centrifuged oocytes were stored at 16° C in ND96 solution for at least 30 min before microinjection of DNA (see below). In one experimental series, the centrifugation force applied ranged between 100 – 3,000 g to characterize the procedure.

### DNA microinjection

The most common dyes used to study molecular permeability through large-pore channels, such as ethidium and YO-PRO, require intercalation with helical polynucleotides to strongly increase their fluorescent emission. To enhance the sensitivity of our technique and to provide a vast excess of binding sites for the polynucleotide-dependent fluorescent molecules, centrifuged oocytes were microinjected with ~50 nL of a solution containing DNA extracted from salmon testes (0.5 mg/mL) and 4’,6-diamidino-2-phenylindole dilactate (DAPI, 100 μM). Microinjection was performed using the Nanoliter 2010 injection system (World Precision Instruments, Sarasota, FL, USA). The DNA-containing pipette was placed in the line that separates the animal pole from the vegetal pole, and microinjections always were oriented toward the translucent zone (animal pole). DAPI was co-injected to visualize the location of the DNA where fluorescence will be emitted after the binding of the permeable molecules tested, in this case, ethidium or YO-PRO. Importantly, DAPI injection did not compromise the sensitivity to other fluorescent dyes, such as ethidium (Supplementary Fig. 1). After microinjection, oocytes were stored in ND96 solution at 16 °C. Resting membrane potential became less negative immediately after centrifugation but soon recovered. Therefore, we kept the oocytes in ND96 for at least 2h before starting the experiments.

### Fluorescence microscopy

Oocytes were transferred to a chamber containing Ringer solution (composition in mM: 115 NaCl, 2 KCl, 1.8 CaCl_2_, 5 HEPES, adjusted to pH = 7.4) and placed on the stage of an inverted epifluorescence microscope (Zeiss Axiovert 200M). Images were acquired using a Zeiss AxioCam MRm camera and the AxioVision 4.8 software (Carl Zeiss, Germany).

### Electrophysiology

Whole-cell recordings and membrane potential measurements were performed using the two-electrode voltage-clamp technique (TEVC). For TEVC, two pulled borosilicate glass micropipettes were filled with 3 M KCl obtaining resistances ranging 0.2 – 2 MΩ. Electrodes for voltage recordings and current injection were mounted on separate micromanipulators (Burleigh PCS-5000, USA) on the stage of a Nikon Eclipse Ti Microscope. The electrical signal was amplified using an Oocyte Clamp Amplifier (OC-725C, Warner Instrument Corp., USA) and digitized by a data acquisition system (Digidata 1440A, Molecular Devices, USA). Data were sampled at 2 kHz, acquired at room temperature (20-22 °C), and analyzed using the pClamp 10 software (Molecular Devices, USA).

### Dye uptake

Oocytes were individually transferred to a perfusion chamber containing Ringer solution and placed on the stage of an inverted epifluorescence microscope (Nikon Eclipse Ti). The chamber was adapted to include a small hole (diameter: 1 mm), designed to keep the oocyte immobile in the center of the microscope platform. A transparent coverslip at the bottom of the hole faced the objective of the microscope. Thus, the oocyte was positioned in the central hole with the vegetal pole facing up and the translucent side facing the objective of the microscope. Then, two electrodes were gently placed into the vegetal pole to perform TEVC in the same oocyte in which dye uptake was evaluated (Fig. 1E). The uptake of 0.1-25 μM YO-PRO (375 Da, charge 2+) or 0.1-50 μM ethidium (314 Da, charge 1+) was assessed at least 10 min after the oocyte was impaled (to allow recovery from the microelectrode impalement).

**Figure 1.**
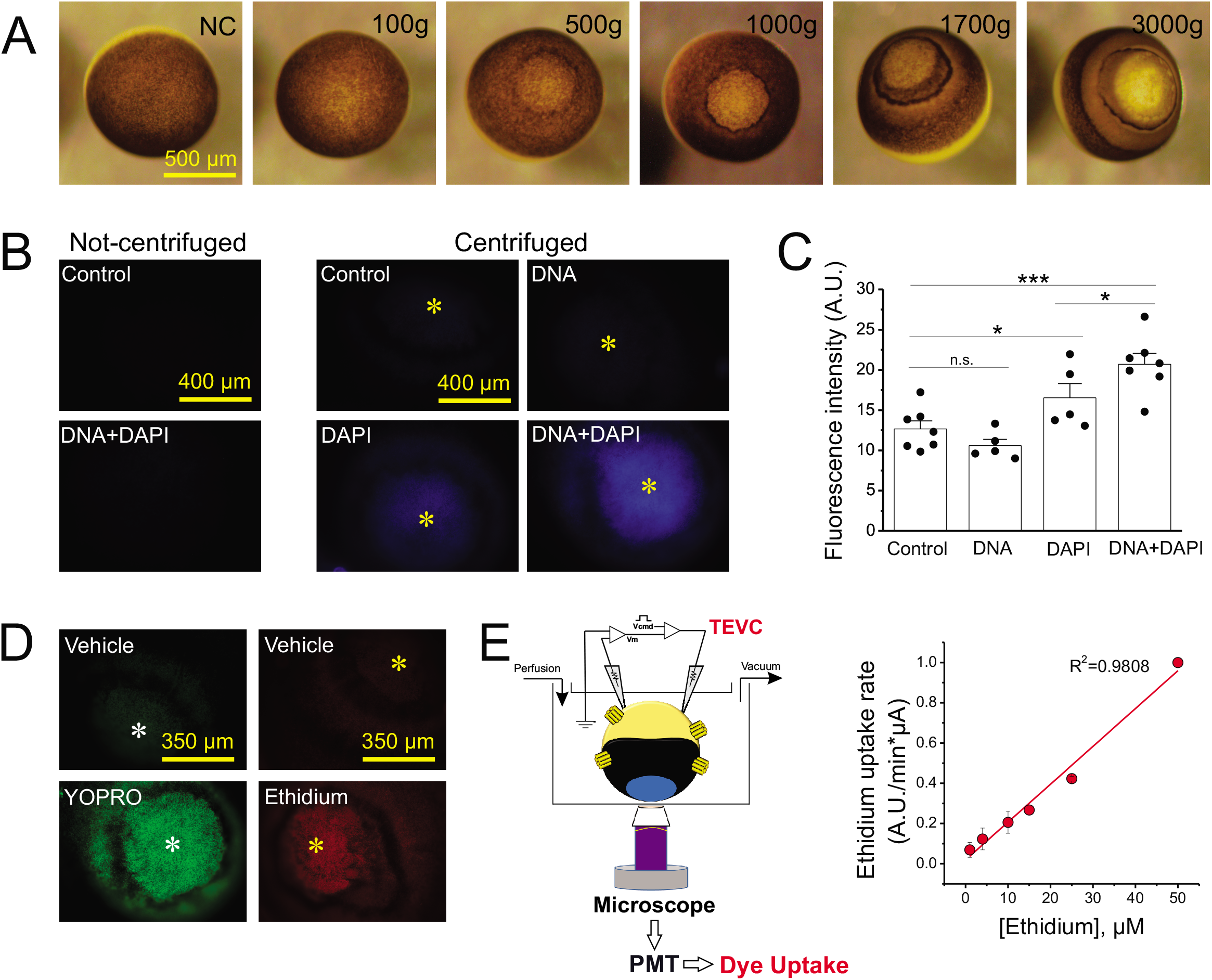
The procedure that turns oocytes translucent improves the visualization of polynucleotide-dependent fluorescent dyes. **(A)** Centrifugal force renders animal pole translucent. **(B)** Representative fluorescent signal (excitation filter: 350 mm /emission filter: 460 nm) detected in non-centrifuged oocytes injected with DNA plus DAPI or water (as control). The fluorescent signal detected in centrifuged oocytes is shown in the right panel. Asterisks indicate the translucent zone in the animal pole. **(C)** Quantification of the fluorescent signal in centrifuged oocytes shown in **B**. **(D)** Representative pictures of YO-PRO or ethidium uptake in centrifuged oocytes. To allow incorporation of dyes into the cytosol, oocytes were permeabilized with 0.01% Triton X-100 and then incubated 3 min with YO-PRO or ethidium (7.5 μM). The vehicle of YO-PRO and ethidium is also shown. **(E)** Left: Scheme of TEVC/Dye uptake assay. Oocytes are centrifuged to create a translucent zone in the animal pole. Then, oocytes are microinjected with 0.5 mg/mL salmon DNA to enhance the detection of polynucleotide-dependent fluorescent dyes. DAPI (100 μM) is coinjected to visualize the injected exogenous DNA. Finally, oocytes are placed in a chamber with the translucent zone facing the objective of an inverted fluorescence microscope. In parallel, the oocyte is impaled following the two-electrode voltage clamp configuration to measure and to control membrane potential. Small fluorescent dyes are incubated or perfused using a gravity-dependent system. Dye uptake is recorded along the time using a photomultiplier tube (PMT). Right panel: Concentration-response curve of ethidium uptake in permeabilized oocytes obtained by using the TEVC/Dye uptake assay. * = p<0.05, *** = p<0.001 by one-way ANOVA plus Newman-Keuls post hoc test.

Dye uptake was recorded in real-time using a photomultiplier tube (Thorlab, PMT1002) connected to a data acquisition system (Digidata 1440A, Molecular Devices, USA). Data were acquired at 2 kHz and analyzed using the pClamp 10 software (Molecular Devices, USA). Because extracellular divalent cations (such as, Ca^2+^) inhibit CALHM1 and connexin hemichannels, dye uptake was evaluated using a divalent cation-free Ringer solution (composition in mM: 115 NaCl, 2 KCl, 5 HEPES, adjusted to pH = 7.4). The effect of extracellular Ca^2+^ (1 mM) on molecular permeation through CALHM1 was also evaluated. For experiments using ethidium, a gravity-dependent flow of 0.5 mL/min was used. For experiments carried out with YO-PRO, no flow was used. All measurements were made at room temperature (20-22 °C).

In some experimental series, dye uptake was assessed in permeabilized oocytes. For this, centrifuged oocytes were incubated in ND96 solution containing 0.01% Triton X-100 for 30 s and subsequently washed with ND96 for 1 min. To obtain representative images of dye uptake, some permeabilized oocytes were incubated for 3 min with 7.5 μM YO-PRO, 7.5 μM ethidium bromide, or its vehicles (as control for autofluorescence). To quantify dye uptake over time, permeabilized oocytes were perfused with 0.1-50 μM ethidium bromide.

### Analysis of channel expression and biophysical properties

To obtain accurate kinetic parameters of permeability, we normalized dye uptake to the activity of functional channels at the cell surface. Thus, before dye uptake recordings, we quantified functional channel expression by the analysis of the current activated by an initial voltage-step pulse from −80 mV to 0 mV for 40 s at nominally zero Ca^2+^ (i.e., in divalent cation-free Ringer solution). In parallel, voltage-activated currents from non-injected oocytes (oocytes which did not express exogenous channels) were also recorded to determine the magnitude of any residual endogenous currents in oocytes from the same batch. Only batches of oocytes with negligible endogenous currents were used.

To analyze the biophysical properties of CALHM1 and Cx26, we analyzed the currents activated by voltage-step pulses from −80 mV to 0 mV (60 s, in divalent cation-free Ringer solution). Additionally, Ca^2+^ concentration-response curves were obtained by analyzing the magnitude of tail currents evoked by the depolarizing pulses, as described previously by Lopez et al. (Lopez et al., 2013). The Ca^2+^ concentration-response relationship was fit to the Hill equation using Origin 9 software (OriginLab Corporation, Northampton, MA, USA). Control experiments indicated that voltage-activated currents in CALHM1 expressing oocytes were inhibited by 30 μM ruthenium red, a CALHM1 blocker, and that voltage-activated current in Cx26 and Cx30 expressing oocytes were inhibited by 30 μM Gd^3+^, a connexin blocker (data not shown).

### Quantification of kinetic parameters of molecular permeation

To calculate the maximum transport (V_max_) and the concentration of the transported molecule (dye) where the half of maximum transport is reached (K_M_), YO-PRO and ethidium uptake rates were fit to a Michaelis-Menten equation using Origin 9 software (OriginLab Corporation, Northampton, MA, USA). Because of the intrinsic variability of channel expression, YO-PRO and ethidium uptake rates were normalized to the magnitude of tail currents assessed in the oocyte before the dye uptake assay (see above).

### Biotinylation and Western blotting

Oocytes expressing Wild-type Cx30 or Cx30^G11R^ were washed out with cold PBS and then incubated with PBS containing 0.5 mg/mL EZ-Link^TM^ NHS-Biotin (Thermo Fisher Scientific) for 10 minutes at 4°C. Then, oocytes were washed out with PBS plus 15 mM glycine for 15 minutes. Oocytes were homogenized in HEN buffer (composition in mM: 250 HEPES, 1 EDTA, and 0.1 neocuproine, adjusted to pH = 7.7) in presence of protease inhibitors (Thermo Fisher Scientific), and then, centrifuged for 10 minutes at 16,000 g. Pellet was resuspended and incubated with streptavidin agarose (Thermo Fisher Scientific) for 30 minutes at 4°C. Samples were then centrifuged at 16,000 g for 2 minutes. The pellet containing biotinylated proteins was washed out with PBS (pH 7.4). Then, samples containing Laemmli buffer were heated at 100°C for 5 minutes to disrupt biotin–streptavidin interaction. The heated samples were then centrifuged for 1 minute at 16,000 g. Subsequently, proteins from the supernatant were separated by 12% SDS-PAGE and transferred onto a PVDF membrane (BioRad, Hercules, CA, USA). The Signal Enhancer HIKARI (Nacalai Tesque, INC, Japan) was used to incubate the primary (Thermo Fisher Scientific, Cat. # 71-2200, 1:1000) and secondary antibody (Pierce, Rockford, IL, USA; 1:5000). SuperSignal® West Femto (Pierce, Rockford, IL, USA) was used to detect the protein bands. Molecular mass was estimated with pre-stained markers (BioRad, Hercules, CA, USA).

### Chemicals

MS-222, collagenase, penicillin-streptomycin, Ficoll, salmon DNA, HEPES, gadolinium, ruthenium red, glycin, neocuproine, and ethidium bromide were purchased from Sigma-Aldrich (St. Louis, MO, USA). DAPI was obtained from Thermo Fisher Scientific (USA). YO-PRO^TM^-1 Iodide was acquired from Invitrogen (USA).

### Statistical analysis

Values are presented as mean ± SEM. Comparisons between groups were made using unpaired Student’s t-test or one-way ANOVA plus Newman-Keuls post hoc test as appropriate. P < 0.05 was considered significant.

## RESULTS

### Centrifugation and DNA microinjection improve the visualization of polynucleotide-dependent fluorescent dyes

To facilitate optical detection of polynucleotide-dependent fluorescent molecules in Xenopus oocytes, we optimized a centrifugation protocol that renders the animal pole of oocytes translucent. Centrifugation created a g force-dependent translucent window in the animal pole by pulling yolk and pigment granules down towards the vegetal pole (Fig. 1A). To confirm that the protocol improves optical visualization of fluorescent dyes, we microinjected salmon DNA plus DAPI into the oocytes. As expected, in non-centrifuged oocytes the fluorescence emitted by DAPI in the intracellular compartment was weak and diffuse (Fig. 1B). In contrast, the DAPI-dependent fluorescent signal was clearer and more intense when it was observed through the translucent zone of the centrifuged oocytes (Fig. 1B-C). Co-injection of salmon DNA significantly increased the intensity of the fluorescent signal emitted by DAPI, indicating that by increasing the DNA binding sites for DAPI the dynamic range for detection of polynucleotide-dependent fluorescent dyes is enhanced (Fig. 1B-C).

Next, we analyzed the uptake of YO-PRO and ethidium, small DNA-intercalator fluorescent molecules. To this end, we used permeabilized oocytes (see Methods) pre-injected with salmon DNA. The incubation of these oocytes in a solution containing YO-PRO or ethidium increased the fluorescent signal observed in the translucent animal pole (Fig. 1D). As expected from previous observations in Xenopus oocytes, we detected autofluorescence at different wavelengths, but it was significantly lower than the fluorescent signal emitted by DAPI, YO-PRO, or ethidium in centrifuged oocytes containing salmon DNA (Fig. 1B-D).

To increase the sensitivity of detection, we analyzed dye uptake with a photomultiplier tube (PMT) instead of a conventional camera. Using this setup, we found that the uptake of ethidium in permeabilized oocytes was linear as a function of concentration (assessed at 10 min of incubation) indicating that ethidium binding to the DNA did not saturate up to 50 μM (Fig. 1E).

To analyze whether the centrifugation procedure and DNA/DAPI microinjection altered membrane permeability and electrochemical gradients, we measured resting membrane potentials in both non-treated and in centrifuged/microinjected oocytes. In this context, the procedure of centrifugation and subsequent DNA injection in Xenopus oocytes did not change the resting membrane potential (measured 2 h after DNA injection), including those oocytes overexpressing large-pore channels formed by CALHM1, Cx26, or Cx30 (Fig. 2A). To determine whether the procedure to make the oocytes translucent affects the gating of heterologously expressed channels, we analyzed the time course of currents activated by voltage in oocytes expressing CALHM1 or Cx26 stimulated with a divalent cation-free solution. In control (non-centrifuged) oocytes, a single voltage pulse from −80 mV to 0 mV led to slow activation currents and triggered tail currents with slow deactivation kinetic characteristics for these channels (Fig. 2B). The ionic currents recorded in oocytes centrifuged and microinjected with DNA had similar activation and deactivation kinetics. (Fig. 2B). In this context, the current detected in control oocytes overlapped the current observed in translucent oocytes, indicating that centrifugation and microinjection of exogenous DNA does not alter the voltage-sensitivity of CALHM1 and Cx26. To test if the procedure of centrifugation and DNA injection affects Ca^2+^ sensitivity of CALHM1 and connexin hemichannels, we then analyzed the activation of these channels at different concentrations of extracellular Ca^2+^. Notably, the voltage-activated currents recorded in control oocytes were similar to those recorded in translucent oocytes (Fig. 2C). Thus, our data indicate that the centrifugation and subsequent DNA microinjection does not modify the Ca^2+^ sensitivity of CALHM1 and connexin hemichannels.

**Figure 2.**
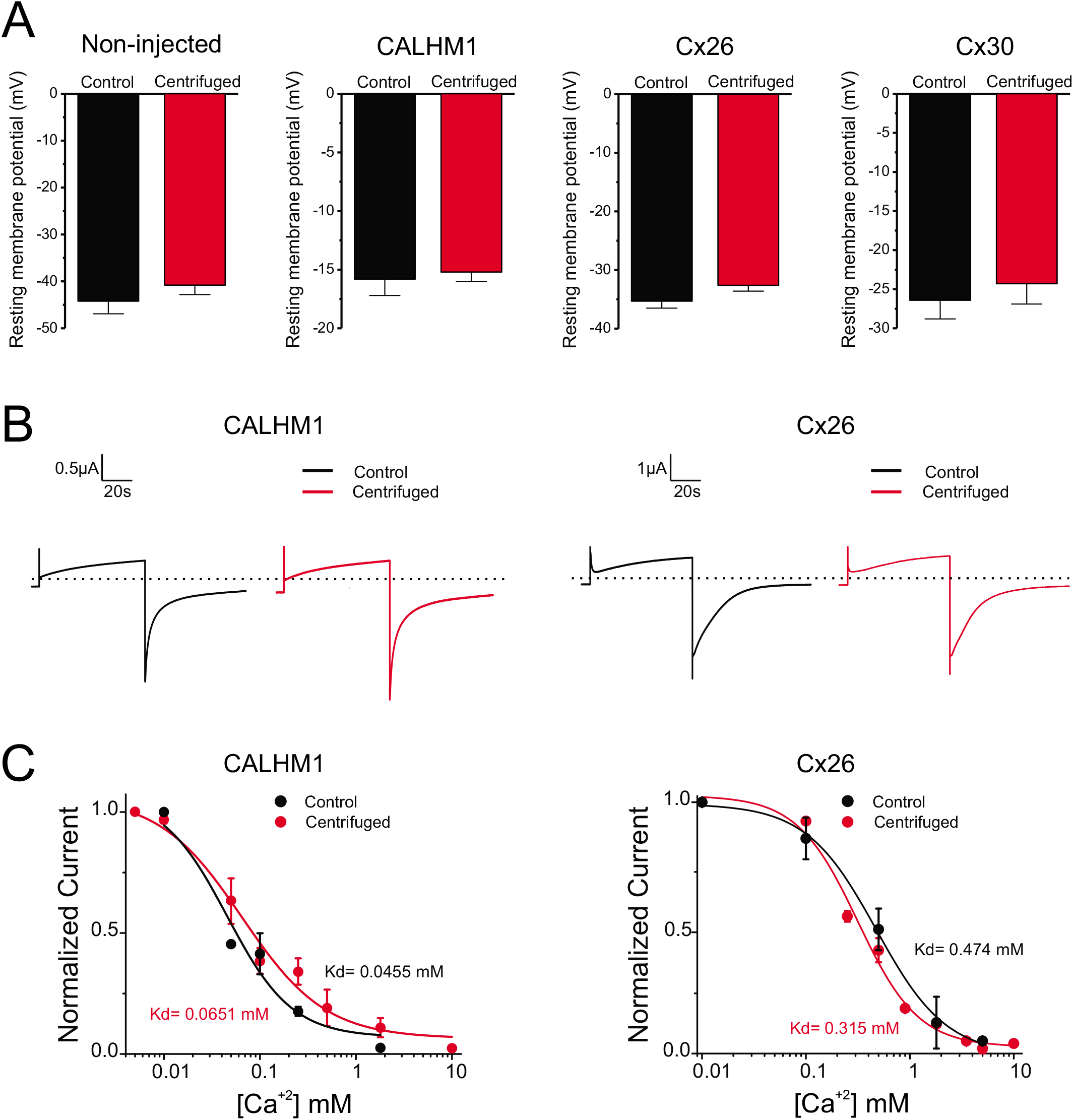
Centrifugation and DNA injection do not affect the biophysical properties of CALHM1 channels and connexin hemichannels. **(A)** Resting membrane potentials recorded in control (non-centrifuged) and centrifuged oocytes. Values were obtained from oocytes perfused with Ringer solution in the presence of 1.8 mM extracellular Ca^2+^. Membrane potentials were measured in non-injected oocytes and in oocytes expressing Cx30, Cx26, and CALHM1. **(B)** Representative current traces elicited by a voltage pulse from −80 to 0 mV in CALHM1 or Cx26 expressing oocytes. Voltage-activated currents were evaluated before and after centrifugation in a divalent cation-free ringer solution. **(C)** Effect of extracellular Ca^2+^ on voltage-activated currents mediated by CALHM1 channels or Cx26 hemichannels.

### The permeation of YO-PRO through the CALHM1 channel is saturable and Ca^2+^- and voltage-dependent

We first studied YO-PRO permeability through CALHM1 channels at resting membrane potential in a divalent cation-free solution. As expected, we did not detect YO-PRO uptake in translucent oocytes that did not express CALHM1 (Fig. 3A, black trace). In contrast, uptake of YO-PRO was clearly evident in oocytes that expressed CALHM1, and the magnitude of dye flux depended on the concentration of YO-PRO (Fig. 3A). The analysis of YO-PRO uptake rates revealed that the permeation of this molecule was saturable (Fig. 3B). The incorporation of 1 mM Ca^2+^, an inhibitor of CALHM1 channels, in the extracellular solution decreased the YO-PRO uptake rate. Theoretically, dye uptake depends on channel expression and channel activity. Consistent with this notion, YO-PRO uptake strongly correlated with the magnitude of CALHM1 channel tail currents (Fig. 3C). To control for variability in channel expression, dye uptake was normalized to the CALHM1 current recorded in the same oocyte. We thereby obtained an accurate and reliable quantification of kinetic parameters for YO-PRO transport, confirming that 1 mM extracellular Ca^2+^ decreased V_max_ by almost half (0.180 ± 0.0063 versus 0.094 ± 0.0056 A.U./min*μA) without affect on the apparent K_M_ (7.50 ± 1.24 versus 8.58 ± 0.24 μM) (Fig. 3D).

**Figure 3.**
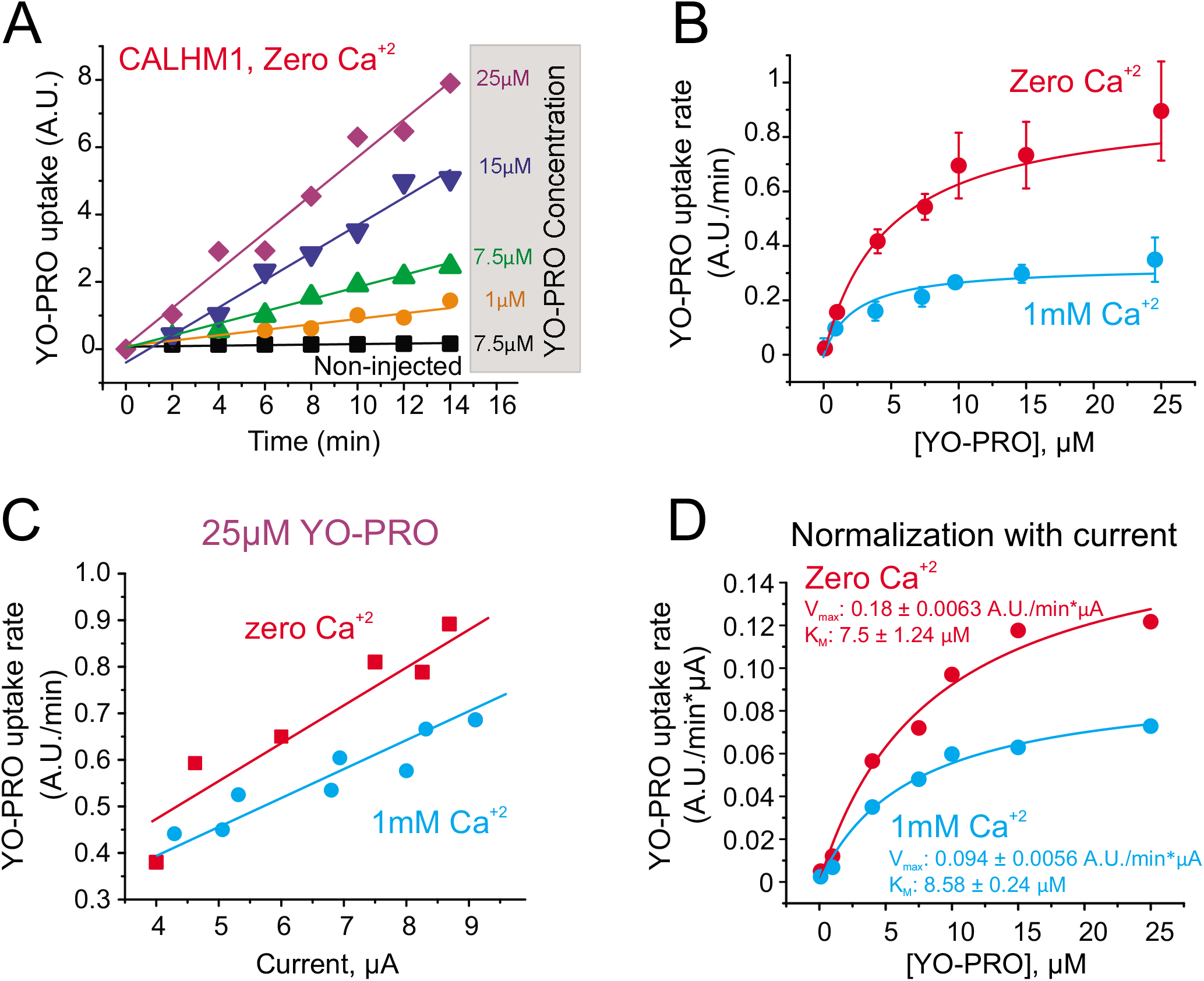
YO-PRO permeation through CALHM1 channels is saturable and Ca^2+^-dependent. **(A)** Representative time course of YO-PRO uptake in non-injected oocytes (black line) or in oocytes expressing CALHM1. Recordings were performed in a divalent cation-free Ringer solution (i.e., zero extracellular Ca^2+^) and at resting membrane potential. **(B)** Analysis of YO-PRO uptake rates in oocytes expressing CALHM1 in divalent cation-free Ringer solution or in the presence of 1 mM extracellular Ca^2+^. **(C)** YO-PRO uptake is correlated to CALHM1 expression (evaluated as the voltage-activated current). **(D)** Kinetic parameters of YO-PRO uptake obtained after normalization by CALHM1 expression (see Methods). Data in D were fit to a Michaelis-Menten equation to obtain V_max_ and K_M_.

Using the TEVC/Dye uptake assay, we explored the voltage-dependency of molecular permeation and atomic ion flux in CALHM1 channels. In the absence of divalent cations, a high conductance was detected at both negative and positive membrane potentials (Fig. 4A). Notably, at 1 mM extracellular Ca^2+^, ionic currents were not detected at potentials more negative than +20 mV (Fig. 4A). At 1 mM extracellular Ca^2+^, YO-PRO uptake progressively increased as membrane potential was clamped at more positive values, despite the fact that electrochemical gradient for YO-PRO is lower at positive membrane potentials (Fig. 4A). Interestingly, YO-PRO uptake was detected at voltages more negative than +20 mV, suggesting that molecular transport occurred even when there is low or no detectable atomic ion permeation. (Fig. 4A). The analysis of the effect of 1 mM extracellular Ca^2+^ on the voltage-activated currents and the maximum flux of YO-PRO revealed that Ca^2+^ inhibited ~95% of ionic currents but only ~48% of YO-PRO uptake (Fig. 4B).

**Figure 4.**
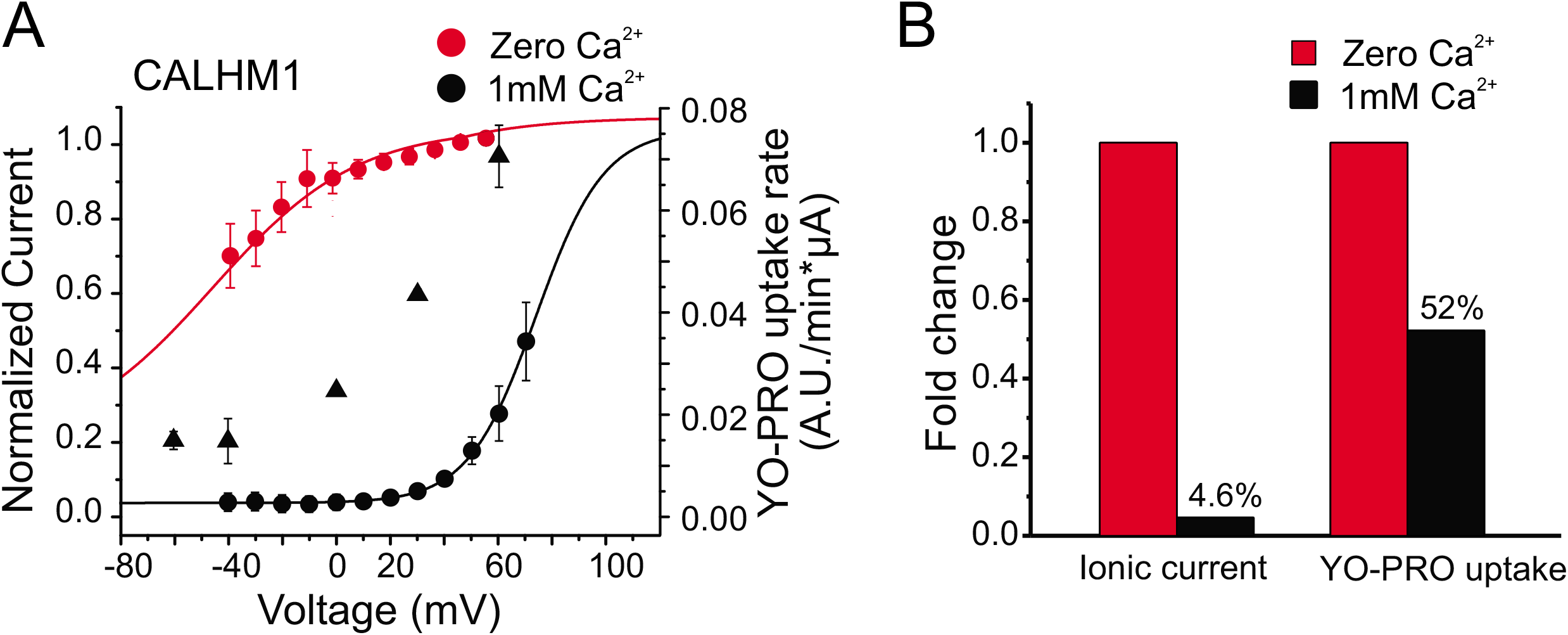
Permeability of molecules and ionic currents in CALHM1 channels can be differentially affected. **(A)** Voltage-dependence of ionic current (circles) and YO-PRO uptake (triangles). The ionic current was evaluated in divalent cation-free Ringer solution or in the presence of 1 mM extracellular Ca^2+^. The current was normalized regarding the current recorded in divalent cation-free Ringer solution. The uptake of 7.5 μM YO-PRO was evaluated at different controlled membrane potentials in the presence of 1 mM extracellular Ca^2+^. **(B)** Analysis of the effect of 1 mM extracellular Ca^2+^ on the ionic currents and the maximum flux of YO-PRO (V_max_).

### Cx30 hemichannels also display saturable transport to small molecules

To evaluate if other large-pore channels display saturating transport to molecules, we analyzed molecular permeability in translucent oocytes expressing connexin hemichannels at resting membrane potentials. We found that Cx30, but not Cx26 was permeable to ethidium (Fig. 5B), consistent with a previous report (Hansen et al., 2014b). Therefore, we decided to evaluate the permeability of ethidium through Cx30 hemichannels. The perfusion of a divalent cation-free solution increased the ethidium uptake rate through Cx30 hemichannels, and the magnitude of the response was dependent on ethidium concentration (Fig. 5A-B). Importantly, the absence of divalent cations did not increase ethidium uptake in translucent oocytes that did not express Cx30 (Fig. 5A-B). The analysis of the ethidium uptake rate revealed that the transport of this molecule through Cx30 hemichannels was saturable at micromolar concentrations (Fig. 5A). Similar to the observations in CALHM1 expressing oocytes, ethidium uptake correlated with the magnitude of tail currents dependent on Cx30 channels (Fig. 5C). The normalization of dye uptake with the current evoked by Cx30 channel opening confirmed that ethidium permeation was saturable with a V_max_ = 0.078 ± 0.015 A.U./min*μA and a K_M_ = 8.10 ± 1.65 μM (Fig. 5D). Overall, our data confirm that Cx30 displays properties of saturating transport similar to that observed for CALHM1 channels.

**Figure 5.**
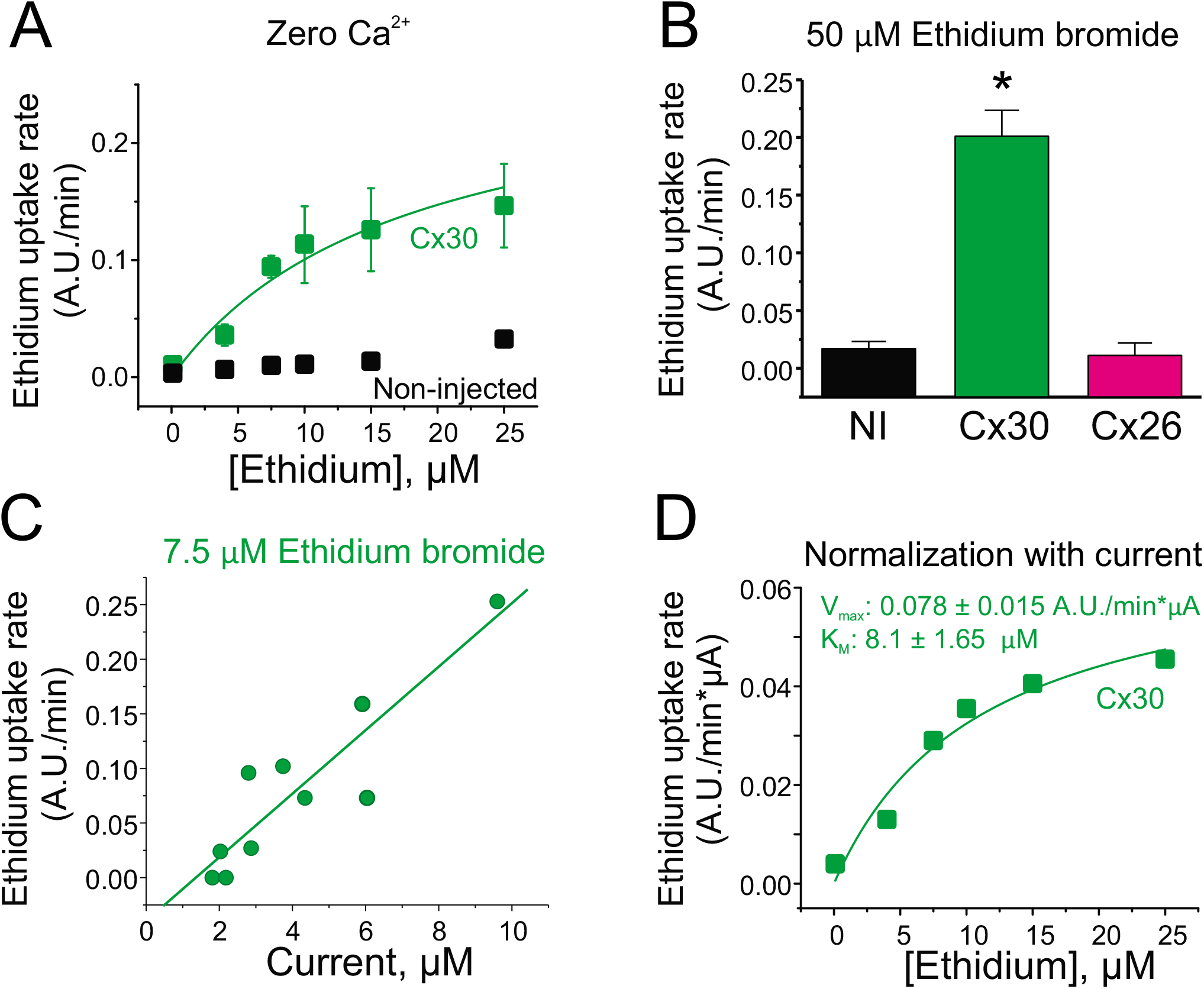
Ethidium permeation through Cx30 hemichannels is saturable and has quantifiable kinetic properties. **(A)** Ethidium uptake obtained in oocytes stimulated with a divalent cation-free Ringer solution at resting membrane potential. Ethidium uptake was evaluated in oocytes expressing Cx30 (green squares) and in non-injected oocytes, which does not express Cx30 (black squares). **(B)** Ethidium uptake rates detected in non-injected oocytes (NI), or in oocytes expressing Cx30 and Cx26. In these experiments, dye uptake was measured in oocytes incubated with a divalent cation-free Ringer solution plus 50 μM ethidium bromide. Ethidium uptake was evaluated at resting membrane potential. **(C)** Ethidium uptake is correlated to Cx30 expression (evaluated as the voltage-activated current). **(D)** Kinetic parameters of ethidium uptake after normalization by Cx30 expression (see Methods). Data were fit to a Michaelis-Menten equation to obtain V_max_ and K_M_. * = p<0.05, by one-way ANOVA plus Newman-Keuls post hoc test.

### Mutations in the N-terminal region of connexin hemichannels affect the selectivity and transport

It has been thought that gain in hemichannel function is associated with an increase in both ionic and molecular permeability. We tested this hypothesis using the TEVC/Dye uptake assay applied to pathogenic human connexin mutations in the N-terminal region, which has been shown to affect gating and permeation.

We first analyzed the human mutation G11R in Cx30, which produces hidrotic ectodermal dysplasia (Essenfelder et al., 2004; Chen et al., 2010). Consistent with the gain of function proposed for this mutation, the voltage-activated currents in oocytes expressing Cx30^G11R^ were higher compared to wild-type Cx30 (Fig. 6A). The increase of voltage-activated currents was not associated with an increase of Cx30^G11R^ channels in the plasma membrane (Supplementary Fig. 2), suggesting that the mutation G11R increases the open probability of the channel. Ethidium uptake in wild-type Cx30 expressing oocytes was detectable at resting membrane potential and correlated with the magnitude of voltage-activated tail currents of Cx30 hemichannels (Fig. 6A). Strikingly, ethidium uptake in Cx30^G11R^ was not detected even when high Cx30^G11R^-mediated currents were detected (Fig. 6A).

**Figure 6.**
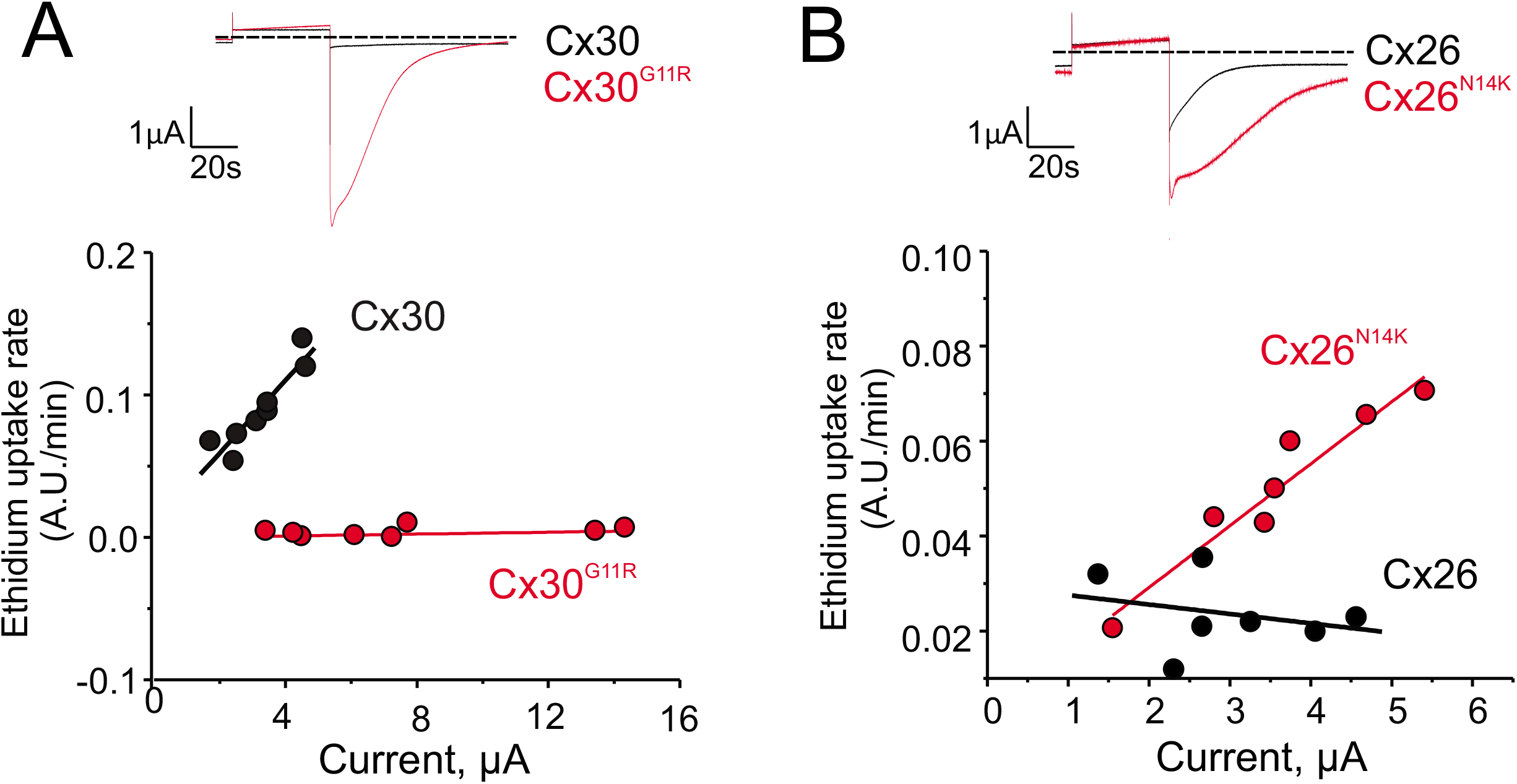
Human mutations in the N-terminal region of connexin hemichannels affect permselectivity. **(A)** Upper panel: Voltage-activated current in oocytes expressing wild-type Cx30 (black trace) or Cx30^G11R^ (red trace). Bottom panel: Ethidium uptake recorded in oocytes with different expression levels of wild-type Cx30 (black circles) or Cx30^G11R^ (red circles). **(B)** Upper panel: Voltage-activated current in oocytes expressing wild-type Cx26 (black trace) or Cx26^N14K^ (red trace). Bottom panel: Ethidium uptake recorded in oocytes with different expression levels of wild-type Cx26 (black circles) or Cx26^N14K^ (red circles). Wild-type Cx30 is permeable to ethidium, but the mutation G11R eliminates permeability to this dye. In contrast, wild-type Cx26 is not permeable to ethidium, but the mutation N14K turns the channel permeable to this molecule. Dye uptake experiments were performed at resting membrane potential, using 7.5 μM ethidium bromide in the bath solution, and at zero extracellular Ca^2+^ (i.e., divalent cation-free Ringer solution).

We then evaluated the human mutation N14K in Cx26, which is also associated with gain in function and produces deafness and skin disorders (Lee et al., 2009; Valdez Capuccino et al., 2019). As expected, the voltage-activated current recorded in oocytes expressing Cx26^N14K^ was higher compared to wild-type Cx26 (Fig. 6B). As seen previously, ethidium uptake in wild-type Cx26 expressing oocytes was negligible at resting membrane potential (Fig. 5B and 6B). In contrast, ethidium uptake was observed in Cx26^N14K^ expressing oocytes, and it was directly proportional to the magnitude of tail currents mediated by Cx26^N14K^ hemichannels (Fig. 6B). Altogether, these data strongly support the notion that atomic ion permeation and molecular permeability are not necessarily coupled in connexin channels, and mutations at the N-terminal region could significantly affect transport of molecules and/or selectivity (i.e., permselectivity).

## DISCUSSION

Large-pore channels such as those formed by connexin, pannexin, and CALHM1 are permeable to both atomic ions and small molecules. Although atomic ion permeation properties through these channels could be successfully assessed using electrophysiological approaches, the study of molecular selectivity and molecular permeation has been limited by qualitative techniques that only allow assessing molecular transport at resting or non-controlled membrane potentials. Here we report a novel technique that allows the study of permselectivity to fluorescent dyes under controlled membrane potentials in Xenopus oocytes. With this methodology, we succeeded in analyzing the voltage-dependence for molecular transport and achieved to quantify accurate kinetic parameters of permeability in CALHM1 channels and Cx30 hemichannels. In addition, we distinguished new biophysical properties that support the hypothesis that molecular permeation is selective, saturable, and it is not coupled to atomic ion fluxes through the pore of large-pore channels.

### Development of a novel technique to analyze molecular transport through large-pore channels

Detection of fluorescence in oocytes is particularly challenging because of the pigments located in the cytosol of the animal pole and the high content of yolk granules that restricts the transmission of fluorescent signals from the intracellular compartment (Dumont, 1972). Therefore, we optimized and characterized a centrifugation-based protocol initially described by Iwao et al. to quantify the cytoplasm volume of non-fertilized oocytes (Iwao et al., 1997). Oocyte centrifugation at 4,000 g was shown to form a translucent cytoplasmic window by pulling down the yolk and pigment granules but affecting cell morphology (Iwao et al., 1997). In our experience, g forces up to 3,000 g created a translucent window in the animal pole without affecting the cell morphology. After the centrifugation step, the subsequent microinjection of DNA and DAPI led, in some cases, to leakage of cytoplasmic content from the sites where microinjections were done. It is important to note that the leakage was observed only in oocytes centrifuged at g forces higher than 1,700 g. The last observation suggests that high centrifugation forces could weaken the plasma membrane. Therefore, to avoid affecting cell morphology, and to retain plasma membrane integrity, we decided to render translucent oocytes using up to 1,700 g for all the TEVC/Dye uptake experiments.

Centrifugation at 1,700 g and subsequent microinjection of DNA did not affect the plasma membrane integrity since resting membrane potential remained unchanged and stable, and oocytes were impermeable to the small charged dyes YO-PRO and ethidium when they did not express exogenous large-pore channels. The presumption that cell function is conserved after the procedure is supported by a previous report showing that the centrifugation of fertilized oocytes at 650 g did not affect their viability and division capability (Iwao et al., 2005). Notably, we observed that viability is not altered at least 2-3 days after centrifugation and DNA microinjection, and most importantly, we demonstrated that this procedure did not affect the biophysical properties of expressed exogenous channels.

Oocytes have a variable intrinsic autofluorescence, which may limit the use of fluorescence to study heterologous expressed ion channels (Lee and Bezanilla, 2019). However, in the translucent oocytes, the fluorescence emitted by the interaction of DAPI, YO-PRO, or ethidium with exogenous salmon DNA in the intracellular compartment was easily distinguished from autofluorescence.

In our TEVC/Dye uptake assay, molecular permeability is assessed by using fluorescent dyes that intercalate with DNA. Therefore, saturation of dye uptake could occur by a reduction of free binding sites in the DNA into the oocytes. We avoided this possibility by injecting extra DNA into the oocyte to enhance the number of binding sites. The co-injection of DNA with DAPI did not affect significantly the binding capacity for ethidium (Supplementary Fig. 1) and most importantly, we observed a linear relationship between fluorescent dye uptake and its concentration in permeabilized cells over the range of concentrations used (Fig. 1E). Thus, we confirmed that binding sites do not saturate in our assay.

We suggest the TEVC/Dye uptake assay could be applicable to almost all large-pore channels. However, a permeable fluorescent dye should be carefully chosen based on the channel under study. Importantly, there is a significant variety of polynucleotide-dependent fluorescent reporters available.

### Analysis of quantitative parameters for molecular transport through large-pore channels

Using the TEVC/dye uptake assay, we found that CALHM1 channels and Cx30 hemichannels showed saturable transport for YO-PRO and ethidium, respectively. Notably, saturation was observed at the low micromolar range, suggesting that these channels behave in some ways like molecule transporters. Interestingly, the quantification of kinetic parameters for the molecular transport in CALHM1 revealed that extracellular Ca^2+^, a well-known regulator of CALHM1 gating did not affect K_M_ but V_max_. These data indicate that Ca^2+^ did not affect the affinity of YO-PRO but maximal molecular flux, probably, as a consequence of the decrease in the open probability.

Because flux is directly related to channel expression (Contreras et al., 2003; Retamal et al., 2007; Schalper et al., 2008; Schalper et al., 2010; Orellana et al., 2011), normalization of dye uptake to the number of functional channels located at the plasma membrane is essential to determine accurate kinetic parameters of permeation. Consequently, the use of fluorescence has been used to estimate the relative channel expression in parallel to functional channel assays (Orellana et al., 2011). The use of fluorescent tags for this purpose, however, has significant limitations. First, it requires tagging the target protein with a fluorescent reporter protein which could affect the gating or permselectivity properties (Bukauskas et al., 2001; Limon et al., 2007; Huang et al., 2011; Gaitán-Peñas et al., 2016). Second, analysis of the fluorescent signal emitted from the plasma membrane is complicated and could be subjective since analysis is performed from one simple focus (Z-axis) and hemichannels need not be uniformly distributed in plasma membrane, as previously reported for CALHM1/3, Cx43, Cx26, and Cx32 (Nakagawa et al., 2011; Orellana et al., 2011; García et al., 2015; Kashio et al., 2019). Third, non-functional or silent channels will be incorporated in the quantification of fluorescence-labeled proteins.

To obtain a quantitative measure of the permeability pathway offered by the channels under study, and to avoid the limitations associated with fluorescent reporters, we determined channel currents by TEVC in the same oocytes in which dye uptake experiments were performed. This current reflects the number of functional channels and their aggregate open probability under each experimental condition, allowing accurate normalization of molecular permeability. To activate CALHM1 channels and connexin hemichannels, we analyzed the magnitude of voltage-activated currents at nominally zero extracellular Ca^2+^, at which a significant increase in the activation of these channels is achieved (Fasciani et al., 2013; Ma et al., 2016). Using this experimental strategy, we observed a linear relationship between molecular transport and the voltage-activated currents, consistent with the notion that macroscopic ionic current is a direct index of the expression of functional plasma membrane channels, as suggested by fluorescent approaches (Contreras et al., 2003).

### Molecular permeability can be assessed at defined membrane potentials

In addition to allow reliable quantification of functional channel expression in parallel with molecular permeability assay, the simultaneous use of TEVC and dye uptake assay have two important advantages for study of permeation of molecules through large-pore channels. First, the whole-cell configuration permits discriminate identification of healthy oocytes (i.e., those with an intact plasma membrane) by the recording of stable currents and resting membrane potentials before and throughout the experiment. Second, the control of the membrane potential allows determination of permselectivity and kinetic parameters in cells at specific/controlled membrane potentials. Thus, the TEVC/dye uptake assay enables the analysis of the voltage-dependency of molecular transport through large-pore channels. Until now, the study of molecular transport through large-pore channels has been performed mostly at resting membrane potentials, at which membrane potential cannot be measured or controlled due to technical limitations. Our methodology will contribute to addressing relevant questions related to the voltage regulation of the transport in large pore channels. This might be particularly important for those channels expressed in excitable cells.

### Implications for the relation between molecular and atomic ion permeation

The conventional view of wide pores is that both atomic ions and molecules diffuse freely through the pore following their electrochemical gradients. In this view, the activation/opening of large-pore channels should proportionally increase the flux of both atomic ions and small molecular permeants as reported for Cx30 hemichannels (Hansen et al., 2014b). However, from the data obtained using the TEVC/dye uptake assay, we suggest that this coupled permeation may not apply to CALHM1 channels. We found that ionic currents through CALHM1 channels are observed only at positive membrane potentials, while significant YO-PRO permeation was detected at negative potentials at which little or no current was detected. Notably, this observation has been previously reported for other large-pore channels. For example, significant Cx43-mediated ethidium permeability was reported at negative resting membrane potential where no atomic ion currents were detected even at the single-channel level (Contreras et al., 2003). Recently, the same phenomenon was observed in pannexin-1 channels (Nielsen et al., 2019a).

The notion that molecular permeation does not directly correlate with, or even be predicted from, atomic ion-dependent currents in large-pore channels is supported by our findings in Cx30 expressing oocytes as well. Cx30 hemichannels are well-known ethidium-permeable channels (Hansen et al., 2014a; Hansen et al., 2014b; Nielsen et al., 2017), but a single mutation in the N-terminal region associated with a considerable gain of atomic ion permeation fully restricted the molecular permeation (Fig. 6A). Consistent with this, alterations in the putative pore residue V37 of Cx30 increase atomic ion flux without altering permeability to ethidium (Nielsen et al., 2017). Interestingly, pharmacological blockers of connexin or pannexin channels can selectively inhibit atomic ion permeation but not molecular uptake (Nielsen et al., 2019a; Nielsen et al., 2019b). Consistent with this, we note that 1 mM extracellular Ca^2+^ reduces ~95% of ionic current but only half of the YO-PRO permeation through CALHM1 channels (Fig. 4B). This evidence supports the hypothesis that large-pore channels are not free diffusion pores for molecules, and distinct mechanisms for atomic ion and molecular transport co-exist (Gaete and Contreras, 2019).

In summary, we report a novel and simple technique that combines electrophysiology and the fluorescence of DNA intercalator dyes to analyze in the same cell the molecular permeability and the atomic ion flux through large-pore channels. The potential use of our TEVC/dye uptake assay could reveal important insights regarding the specific mechanisms of molecular and atomic ion permeability in health and disease.

## ACKNOWLEDGMENTS

This work was supported by the National Institutes of Health/National Institute of General Medical Sciences (Grants RO1-GM099490 to J.E. Contreras and RO1-GM101950 to A.L. Harris and J.E. Contreras), and an American Heart Association (AHA) postdoctoral fellowship 18POST339610107 to M.A. Lillo.

## AUTHOR CONTRIBUTIONS

JEC and PSG designed the research. YL performed molecular biology. WL, MAL and PSG performed research. MAL, PSG, and WL analyzed experimental data. PSG, ALH, and JEC wrote and edited the manuscript.

