## Supplementary material for "A novel voltage clamp/dye uptake assay reveals saturable transport of molecules through CALHM1 and connexin channels"

**Supplementary figure 1.** DAPI does not saturate the number of binding sites available for ethidium intercalation. **(A)** Left panel: Representative recordings of fluorescence emitted by a drop (100 µL) of water (Control, black trace), 1 µM ethidium bromide (red trace), 1 µM ethidium bromide + 0.5 mg/mL salmon DNA (magenta trace), or 1 µM ethidium bromide + 0.5 mg/mL salmon DNA + 100 µM DAPI (blue trace). Fluorescence was detected by a photomultiplier tube (PMT) in a dark room. The horizontal arrow indicates the baseline where the LED is turned off. The horizontal bar indicates the period when samples are excited (3 s, excitation filter: 530 nm; emission filter: 600 nm). Right panel: Analysis of the fluorescence shown in the left panel. **(B)** Fluorescence detected in a microplate reader in a fluorescence spectrophotometer (Cary, Eclipse). Plates containing 0.5 mg/mL salmon DNA were evaluated in the absence and presence of 100 µM DAPI. Fluorescence was analyzed before and after the addition of 1 µM ethidium bromide.

**Supplementary figure 2.** The gain of function of $Cx30^{G11R}$ channels is not associated with an increase in protein expression. The figure shows a representative Western blot of Cx30 expression in Xenopus oocytes. Expression was evaluated 24 and 48 h after cRNA microinjection. In these experiments, oocytes were microinjected with the same amount of cRNA for Wild-type (WT) and $Cx30^{G11R}$. In addition to the total Cx30, we examined the protein located in the plasma membrane by biotinylation (see Methods).

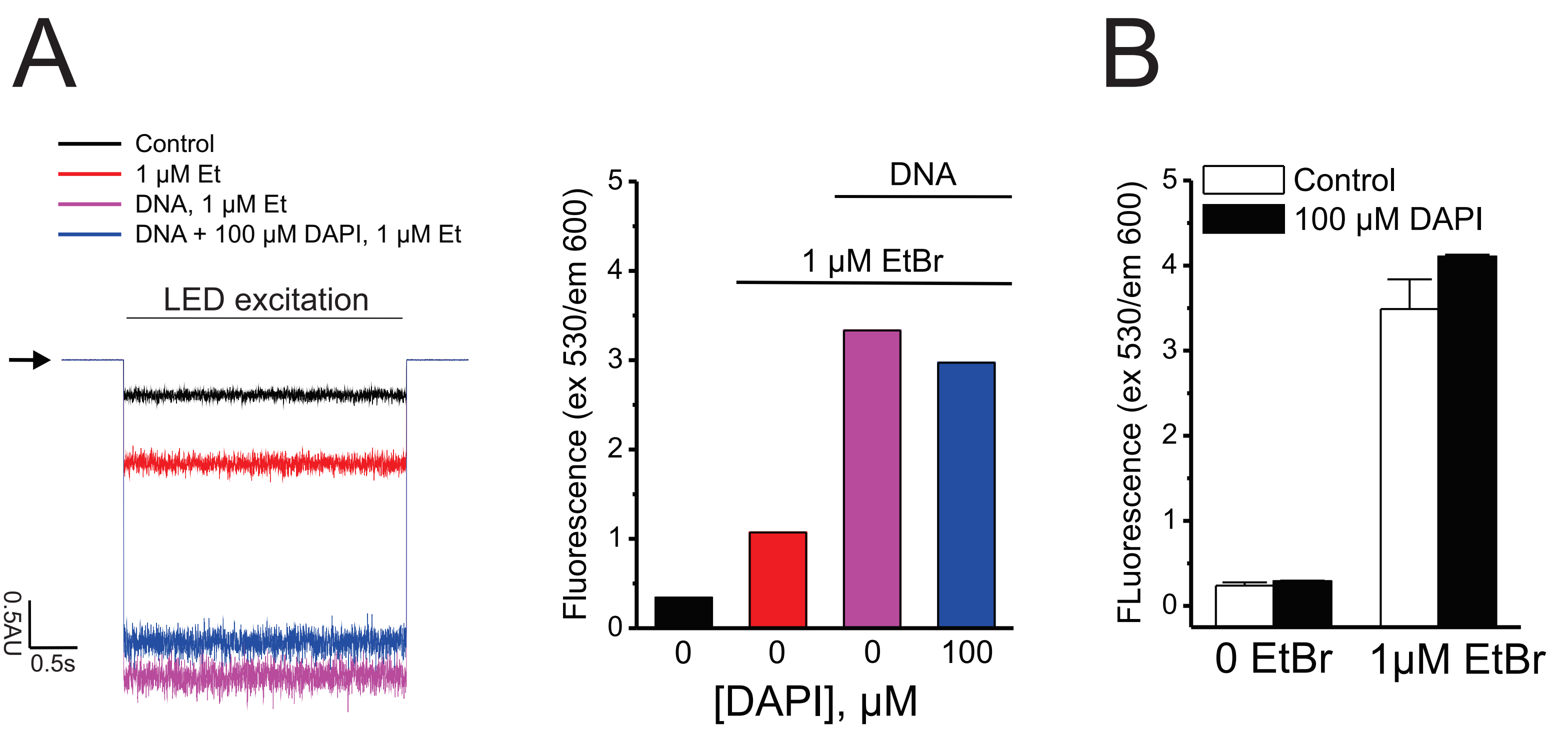


Supplementary Figure 1

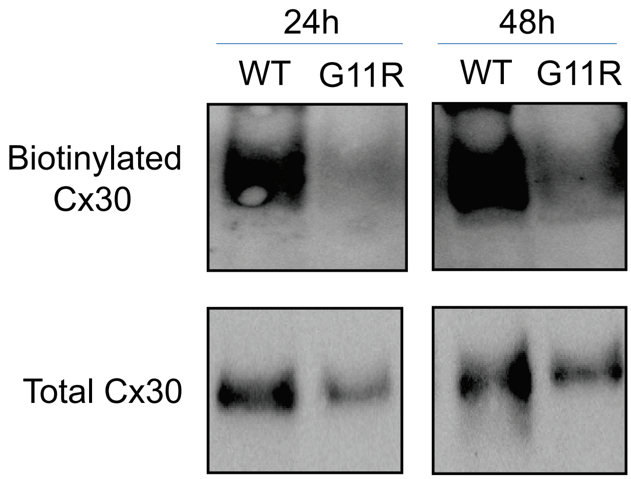


Supplementary Figure 2
